# High-throughput yeast two-hybrid library screening using next generation sequencing

**DOI:** 10.1101/368704

**Authors:** Marie-Laure Erffelinck, Bianca Ribeiro, Maria Perassolo, Laurens Pauwels, Jacob Pollier, Veronique Storme, Alain Goossens

**Affiliations:** Ghent University, Department of Plant Biotechnology and Bioinformatics, Technologiepark 927, B-9052 Ghent, Belgium; VIB Center for Plant Systems Biology, Technologiepark 927, B-9052 Ghent, Belgium; Universidad de Buenos Aires, Facultad de Farmacia y Bioquímica, Departamento de Microbiología, Inmunología y Biotecnología, Cátedra de Biotecnología, Buenos Aires, Argentina; CONICET-Universidad de Buenos Aires, Instituto de Nanobiotecnología (NANOBIOTEC), Buenos Aires, Argentina

## Abstract

Yeast two-hybrid (Y2H) is a well-established genetics-based system that uses yeast to selectively display binary protein-protein interactions (PPIs). To meet the current need to unravel complex PPI networks, several adaptations have been made to establish medium- to high-throughput Y2H screening platforms, with several having successfully incorporated the use of the next-generation sequencing (NGS) technology to increase the scale and sensitivity of the method. However, these have been to date mainly restricted to the use of fully annotated custom-made open reading frame (ORF) libraries and subject to complex downstream data processing. Here, a streamlined high-throughput Y2H library screening strategy, based on integration of Y2H with NGS, called Y2H-seq, was developed, which allows efficient and reliable screening of Y2H cDNA libraries. To generate proof of concept, the method was applied to screen for interaction partners of two key components of the jasmonate signaling machinery in the model plant *Arabidopsis thaliana*, resulting in the identification of several previously reported as well as hitherto unknown interactors. Our Y2H-seq method offers a user-friendly, specific and sensitive screening method that allows high-throughput identification of PPIs without prior knowledge of the organism’s ORFs, thereby extending the method to organisms of which the genome has not entirely been annotated yet. The quantitative NGS readout and the incorporation of background controls allow to increase genome coverage and ultimately dispose of recurrent false positives, thereby overcoming some of the bottlenecks of current Y2H technologies, which will further strengthen the value of the Y2H technology as a discovery platform.

## Introduction

Disentangling protein-protein interaction (PPI) networks is crucial for our understanding of cellular organization and function. To achieve this, a wide range of technologies to identify PPIs has been developed over the last decade [1,2]. One of the most advanced and commonly used methods to identify PPIs *in vivo* under near-physiological conditions is affinity purification coupled to mass spectrometry (AP-MS) [3–5]. Equivalent comprehensive assays to specifically identify binary PPIs include protein domain microarrays and *in vivo* protein fragment complementation assays (PCAs) [6–10]. The principle of PCA is based on the fusion of two hypothetically interacting proteins (bait and prey) to two fragments of a reporter protein. Interaction between the bait and prey proteins results in the reassembly of the reporter protein, followed by its activation. The signal readout can be bioluminescence, fluorescence or cell survival. In the popular yeast two-hybrid (Y2H) method, the bait protein is fused to the DNA binding domain (DBD) and the prey (or prey library in the case of a comprehensive Y2H screening) is fused to the activation domain (AD) of a transcription factor (TF) [11]. Upon association of the hypothetical interactors, the TF is functionally reconstituted and drives the expression of a reporter gene that can be scored by selective growth. Typically, conventional medium-throughput Y2H library screenings are subject to laborious one-by-one clonal identification of interaction partners, but today, proteome-wide mapping of PPIs demands a high-throughput approach. This led for instance to the development of a matrix-based Y2H method that bypassed the inefficient identification by DNA sequencing [12]. Collections of bait and prey strains were automatically combined and arrayed on fixed matrix positions and PPIs were scored as visual readouts. A major drawback of this strategy is the need for pre-assembled libraries based on defined gene models and expensive robotics that are not accessible to every researcher.

Clonal identification of Y2H screening with DNA sequencing has a tremendous negative effect on the efficiency, cost and labor of the method. Furthermore, given the labor-penalty involved with increasing transformation titers, the clonal identification of Y2H interactions is usually not compatible with quantitative assessment of PPI abundances. Therefore, replacing the conventional Y2H screening strategy with a pool-based selection and global identification by NGS, can have three major implications: (i) cost reduction by high-capacity sequencing, (ii) higher sensitivity and (iii) quantification of the abundance of bait-specific interactions. The lab of Marc Vidal pioneered the implementation of the NGS technology for massive parallel Y2H screening in the Stitch-Seq method, mainly to map the human interactome. Herein, single amplicons, concatenating sequences of potentially interacting proteins, serve as template for NGS [13]. Nonetheless, this method remains laborious because it requires clonal isolation and several PCR rounds for PPI identification for each selected colony. The lab of Ulrich Stelzl developed the Y2H-seq method, thereby illustrating the advantage of NGS for Y2H towards scalability by mapping the protein methylation interactome [14]. In this strategy, the use of barcode indexing enables simultaneous sequencing of interacting preys of multiple separate baits in a single Illumina run. This strategy is based on mixing bait and prey pools prior mating, followed by selective growth, and deep-sequencing, but still requires a post-screen binary testing of interacting baits with each of the identified preys. The use of barcodes was further exploited in the Barcode Fusion Genetics-Yeast Two-Hybrid (BFG-Y2H) method. This matrix-Y2H strategy uses Cre-recombinase to create intracellular chimeric barcodes that are derived from protein pairs, thereby enabling immediate identification and quantification of each interaction pair through NGS [15]. Prior to screening and NGS, isolation and sequencing of each barcoded bait and prey clone are essential to associate barcodes to ORFs, which may pose a cost restriction for massive screening purposes. The latter was addressed in CrY2H-seq, which introduced a Cre-recombinase interaction reporter that endorses fusion of the coding sequences of two interacting proteins, followed by NGS to identify these interactions *en masse* [16]. The latter method was employed to uncover the transcription factor interactome of *A. thaliana*.

All of the above-mentioned Y2H-NGS strategies focus on increased capacity, efficiency and sensitivity, although they may face some lack in specificity or do not fully exploit the quantification potential of NGS coupled to Y2H. Furthermore, construction of full-length ORF libraries are necessary, thereby restricting these methods to organisms of which the genomes are well annotated or to ‘defined’ gene models, which for instance cannot take alternative splicing, alternative start codon use or transcript processing into account.

Here, we discuss a user-friendly and standardized Y2H-NGS workflow (‘Y2H-seq’), complementary to the matrix-Y2H approaches, which allows rapid identification of interaction partners of a bait of interest in the organism of choice without the need for expensive robotics. The Y2H-seq screening method generates a quantitative readout that, through the use of control screens, allows to eliminate false-positive PPIs to boost the specificity of the method and thereby avoiding unnecessary downstream experimental binary interaction verification. Furthermore, the method is not dependent on predefined and prefabricated ORF libraries but on cDNA libraries, and is therefore principally applicable to every organism regardless of the annotation status of its genome. The functionality of our methodology is validated here by implementing it on two well-studied members of the jasmonate (JA) signaling cascade in the model plant *Arabidopsis thaliana*, i.e. TOPLESS (TPL) and Novel Interactor of JAZ (NINJA), respectively encoded by the loci AT1G15750 and AT4G28910 [17–25].

## Material and methods

### Gene Cloning

All cloning was carried out by Gateway^®^ recombination (Thermo Fisher Scientific, Waltham, MA, USA). The full-length coding sequence of *IAA17* was PCR-amplified (for primers, see S2 Table) and recombined in the donor vector pDONR221. All other entry clones had previously been generated [17,26].

### Binary Y2H analysis

Y2H analysis was performed as described [27] using the GAL4 system [27], in which bait and prey were fused to the GAL4-AD or GAL4-BD via cloning into pDEST™22 or pDEST™32, respectively. The *Saccharomyces cerevisiae* PJ69-4α yeast strain [28] was co-transformed with bait and prey constructs using the polyethylene glycol (PEG)/lithium acetate method. Transformants were selected on SD medium lacking Leu and Trp (Clontech, France). Three individual colonies were grown overnight in liquid cultures at 30°C and 10- or 100-fold dilutions were dropped on control (SD-Leu-Trp) and selective media (SD-Leu-Trp-His).

### Y2H screening

Yeast transformation was performed as described by Cuéllar-Pérez *et al.*, (2013) [27]. The *S. cerevisiae* PJ69-4α yeast strain was transformed in two transformation rounds, respectively with 0.5 μg of bait plasmid DNA and 50 μg of cDNA prey library plasmid DNA using the PEG/lithium acetate method. At least 10^6^ transformants were plated on control (SD-Leu-Trp) and selective media lacking Leu, Trp and His supplemented with 5 mM 3-AT (Sigma-Aldrich, Saint Louis, MO, USA).

### Y2H cDNA library used to perform the Y2H screening

The ProQuest two-hybrid cDNA library was generated by cDNA synthesis from RNA extracted from *A. thaliana* suspension cells AT7, cloned into pEXP-AD502 vector (ProQuest), equivalent to pDEST™22 vector (Thermo Fisher Scientific) and electroporated in the DH10B-Ton A (T1 and T5 phage resistance) cells (Thermo Fisher Scientific). The average insert size was 1.1 kb and the number of primary clones was 5.3 x 10^6^ cfu with a 100% insert coverage.

### Sanger sequencing

A minimum of ten random colonies of the Y2H screening plates were streaked out on solid SD-Leu-Trp-His selective medium with 5mM 3-AT (Sigma-Aldrich, Saint Louis, MO, USA) and incubated for 48 h at 30°C. Each streaked out colony was inoculated in liquid SD-Leu-Trp-His selective medium and incubated overnight at 30°C at 230 rpm. Subsequent yeast plasmid isolation was carried out using the Zymoprep™ Yeast Plasmid Miniprep I Kit (Zymo Research, Irvine, CA, USA) according to the manufacturer’s instructions. The cDNA inserts of the prey plasmids (pDEST™22-insert) were PCR-amplified using backbone-specific primers (S2 Table) and Sanger-sequenced.

### Semi-quantitative qPCR

Colonies of the Y2H screening plates were dissolved and pooled in 10-15 mL of ultrapure water and plasmids were collected using the Zymoprep™ Yeast Plasmid Miniprep II kit (Zymo Research, Irvine, CA, USA). Prey constructs were amplified via PCR using Q5^®^ High-Fidelity DNA Polymerase (New England Biolabs, Ipswich, MA, USA) and generic pDEST™22 primers that bind to the GAL4AD and the region flanking the attR1 site (S2 Table). The following program was used: initial denaturation (98°C, 30 s), 35 amplification cycles (denaturation 98°C, 10 s; annealing 55°C, 30 s; elongation 72°C, 2.5 min), final extension (72°C, 5 min). The PCR mixture was purified using the CleanPCR kit (CleanNA, Alphen aan den Rijn, The Netherlands) and 40 ng of the purified PCR product was used for semi-quantitative qPCRs, which were carried out with a Lightcycler 480 (Roche Diagnostics, Brussels, Belgium) and the Lightcycler 480 SYBR Green I Master kit (Roche). Specific primers (S2 Table) and GoTaq^®^ DNA polymerase (Promega, Fitchburg, WI, USA) were used for amplification of 40 ng of purified PCR product with the following program: initial denaturation (95°C, 5 min), 40 amplification cycles (denaturation 95°C, 30 s; annealing 60°C, 30 s; elongation 72°C, 60 s), final extension (72°C, 5 min). As a reference, a short sequence originating from the AD of pDEST™22 was used. For the relative quantification with the reference gene, qBase was used [29].

### NGS data processing

The samples were sequenced by Illumina HiSeq 2000 125-bp paired-end reads. Data mapping and filtering were carried out through an in-house generated pipeline. To avoid sequencing artifacts such as read errors, primers, adapter and vector sequence contamination and PCR bias, a quality check was performed on the raw sequencing data. The quality control and trimming were performed with Trimmomatic [30]. Subsequently, the processed sequencing reads were mapped against the *Arabidopsis* reference genome, downloaded from TAIR (The Arabidopsis Information Resource, http://arabidopsis.org), by TopHat [31], which uses the Bowtie program as an alignment engine. In addition, TopHat requires SAM (Sequence Alignment/Map) tools to be installed. The cufflinks program was used to count the expression of each gene and report it as raw reads and FPKM. To determine possible interactors, following steps were taken. Genes with less than six read counts were not considered. Zero counts in the negative control sample were replaced by 1 to avoid division by 0. These genes were flagged to keep track of these imputations. FPKM values were calculated for each gene in both the sample and the negative control.

Subsequently, the SNR was calculated for each gene as the ratio of the sample FPKM value to the negative control FPKM value. Genes with an SNR_NINJA/EMPTY_ or SNR_TPL-N/EMPTY_ higher than the arbitrary threshold of 11, were considered to be potential interaction partners of the bait gene.

## Results

### Selection of baits

JAs are phytohormones that regulate the plant’s defense and modulate several developmental processes. The production of JAs via the oxylipin biosynthetic pathway leads to the accumulation of bioactive (+)-7-*iso*-jasmonoyl-L-isoleucine (JA-Ile). JA-Ile functions as a ligand between the F-box protein coronatine insensitive 1 (COI1) and the JA-ZIM (JAZ) repressor proteins, thereby promoting ubiquitination and subsequent proteasomal degradation of the JAZ proteins [32,33]. Together with the TIFY8, peapod (PPD) and ZIM proteins, the JAZ proteins belong to the TIFY super-family [32, 34–37]. A key regulator in JA signaling in *A. thaliana* is the basic helix-loop-helix (bHLH) TF MYC2, encoded by the locus AT1G32640 [38,39]. In the absence of JA-Ile, MYC2 can physically interact with the JAZ proteins via the Jas motif, which in turn recruit the transcriptional repressor TPL and TPL-related proteins (TRPs) through the adaptor protein NINJA [17]. NINJA acts as a transcriptional repressor that harbors an intrinsic TPL-binding ETHYLENE RESPONSE FACTOR (ERF)-associated amphiphilic repression (EAR) motif mediating its activity (Fig 1) [17]. NINJA can also interact with non-JAZ TIFY proteins, demonstrating its role in processes other than JA signaling [17,34,37,40,41]. Likewise, TPL is associated with various cellular processes through its capacity to interact with a compendium of diverse proteins [17–24, 37]. For instance, TPL can bind to PEAPOD proteins through the adapter proteins KIX8 and KIX9 to negatively regulate meristemoidal division in *A. thaliana* [37]. A role for TPL modulating brassinazole resistant 1 (BZR1)-regulated cell elongation and brassinosteroid-mediated control of shoot boundaries and root meristem development through interaction with the TF bri1-ems-suppressor 1 (BES1) has been described [22,24]. TPL can also be recruited by CC-type glutaredoxins to target TGA-dependent promoters to control development- and stress-associated processes.

**Figure 1.**
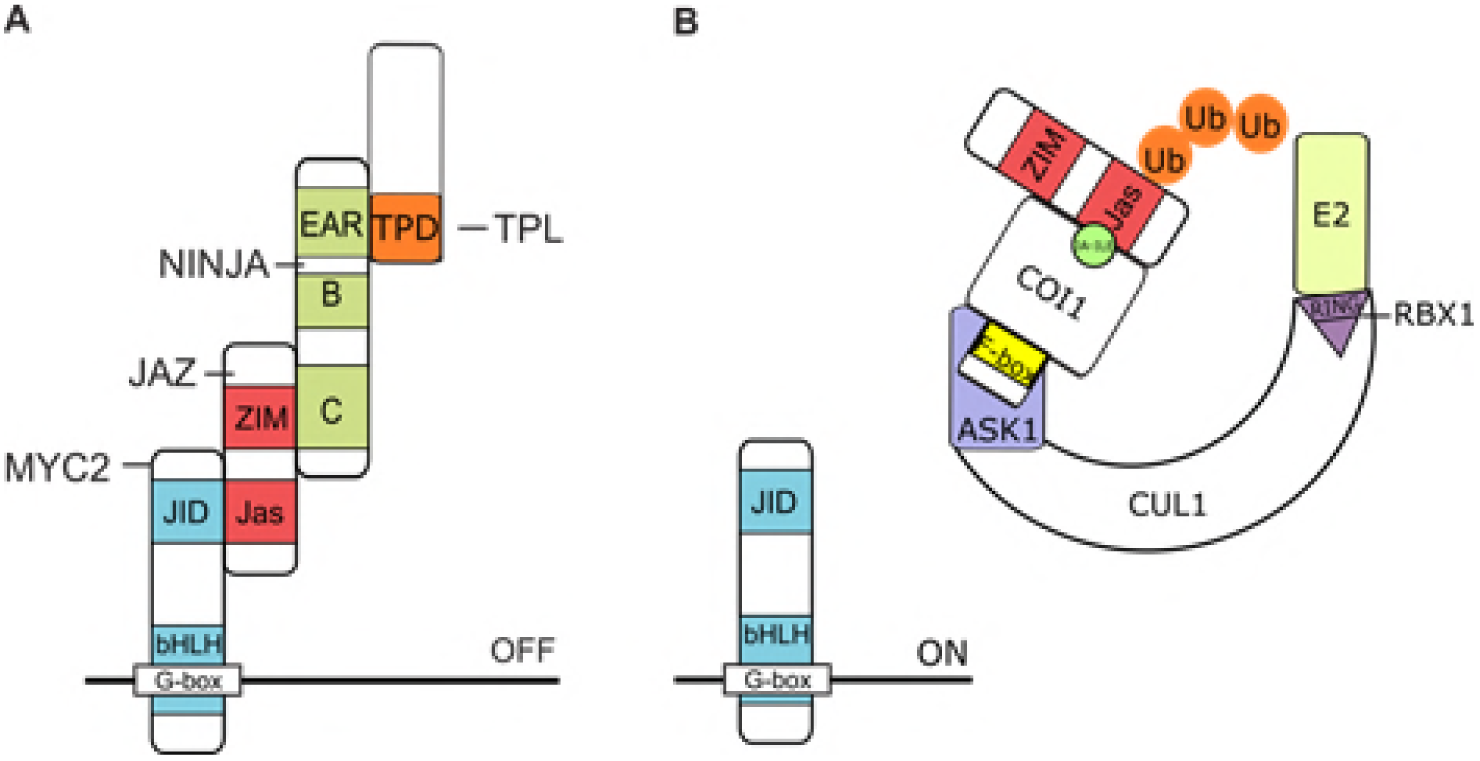
Function of TOPLESS and NINJA in JA signaling in *A. thaliana*. (A) In the absence of JAs, bHLH-type MYC TFs interact with the Jas domain of JAZ proteins that in turn interact with NINJA via their ZIM domain. The EAR motif of NINJA is essential for recruitment of the TPL co-repressors through the TPL domain (TPD). (B) In the presence of JA-Ile, JAZ proteins interact with the ubiquitin E3 ligase SCF^COI1^ complex, leading to the proteasomal degradation of JAZs and consequent release of the NINJA–TPL complex from the MYC TFs, which leads to the transcriptional activation of JA- responsive genes by de-repressed MYC TFs.

Because various direct interactors have been described for both NINJA and TPL proteins and because these are currently still heavily investigated for potential novel roles and links with different signaling pathways and cellular processes, NINJA and TPL were chosen as ideal bait proteins to develop, establish and validate our Y2H-seq methodology. Notably, whereas we used the full-length ORF of NINJA as a bait, for TPL only the amino-terminal region (AA 1- 188; TPL-N) was used as a bait because this domain contains the lissencephaly homologous (LisH) dimerization and C-terminal to LisH (CTLH) motifs, which are together required and sufficient for interaction with transcriptional repressors through their EAR motif.

### The Y2H-seq flow-chart

An illustration of the general workflow of our Y2H-Seq strategy is given in Figure 2. As indicated above NINJA and TPL-N were used as baits and a Y2H cDNA library originating from *A. thaliana* AT7 suspension cells was used as prey.

**Figure 2.**
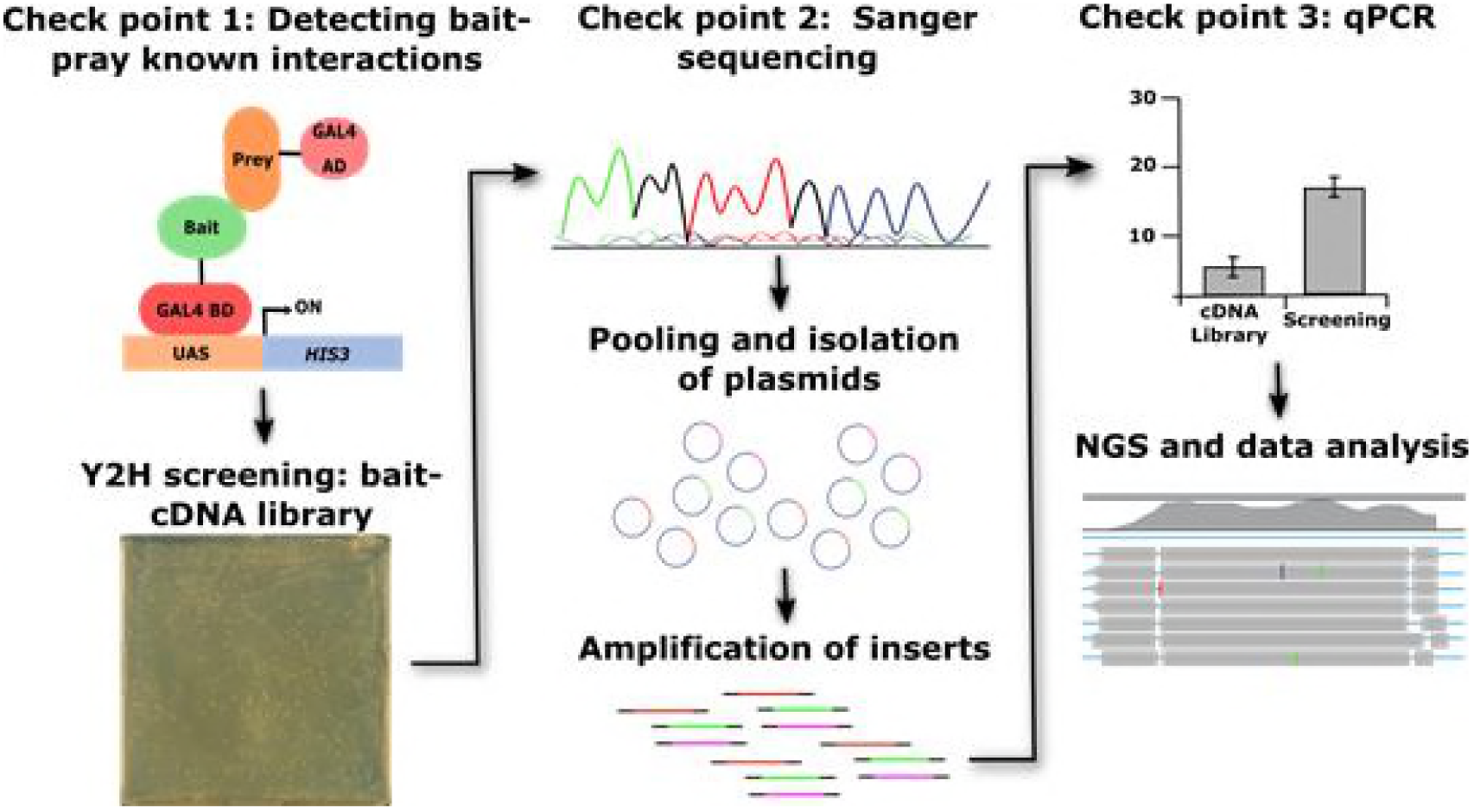
Y2H-seq workflow.

After transformation of the Y2H reporter strain PJ69-4α with the bait plasmids, a first checkpoint is introduced, in which the bait strains were individually co-transformed with positive and negative control prey expression clones to verify functional expression of the baits, to exclude possible auto-activation and to corroborate binding with previously reported interaction partners (Fig 1). Next, the bait strains were used for Y2H-seq screening with the *A. thaliana* Y2H cDNA prey library. Simultaneously, a control screening was performed with the empty expression vector, which will hereafter be referred to as EMPTY. Subsequent to five days of selective growth of the transformed yeast cells, the prey cDNA inserts of about ten individual yeast colonies per screen were Sanger-sequenced (Fig 1). This second checkpoint allowed us to confirm the retrieval of reported interactors as preys. Subsequently, all yeast colonies that survived selective growth were pooled per screen and the cDNA inserts of the prey plasmid pools were amplified by PCR. A third checkpoint consisted of a qPCR analysis with specific primers for genes corresponding to known bait interactors, which allows to assess the representation of known interactors in both screens in a quantitative manner (Fig 1). Prey abundance was quantified relative to that in the *A. thaliana* Y2H cDNA library.

Upon complying the expectations of all three checkpoints, the amplicons of the pooled prey cDNA inserts were sequenced by NGS (Illumina HiSeq 2000 sequencing, 125-bp paired-end reads). The NGS-output was analyzed by an adapted RNA-Seq data processing pipeline, providing a quantitative selection of known and potentially new interactors of NINJA and TPL-N, using the EMPTY screen as control to eliminate false-positive interactions and to correct for the abundance of each prey represented by the Y2H cDNA library.

### Y2H-seq checkpoints

#### Checkpoint 1: exploring auto-activation and functionality of the bait strains

The bait strains were individually co-transformed with positive and negative control preys (Fig 3 and S1 Table) to determine the level of auto-activation of the bait strain and to check whether the bait protein is functionally expressed and consequently can bind previously reported interaction partners [17,25,37].

**Figure 3.**
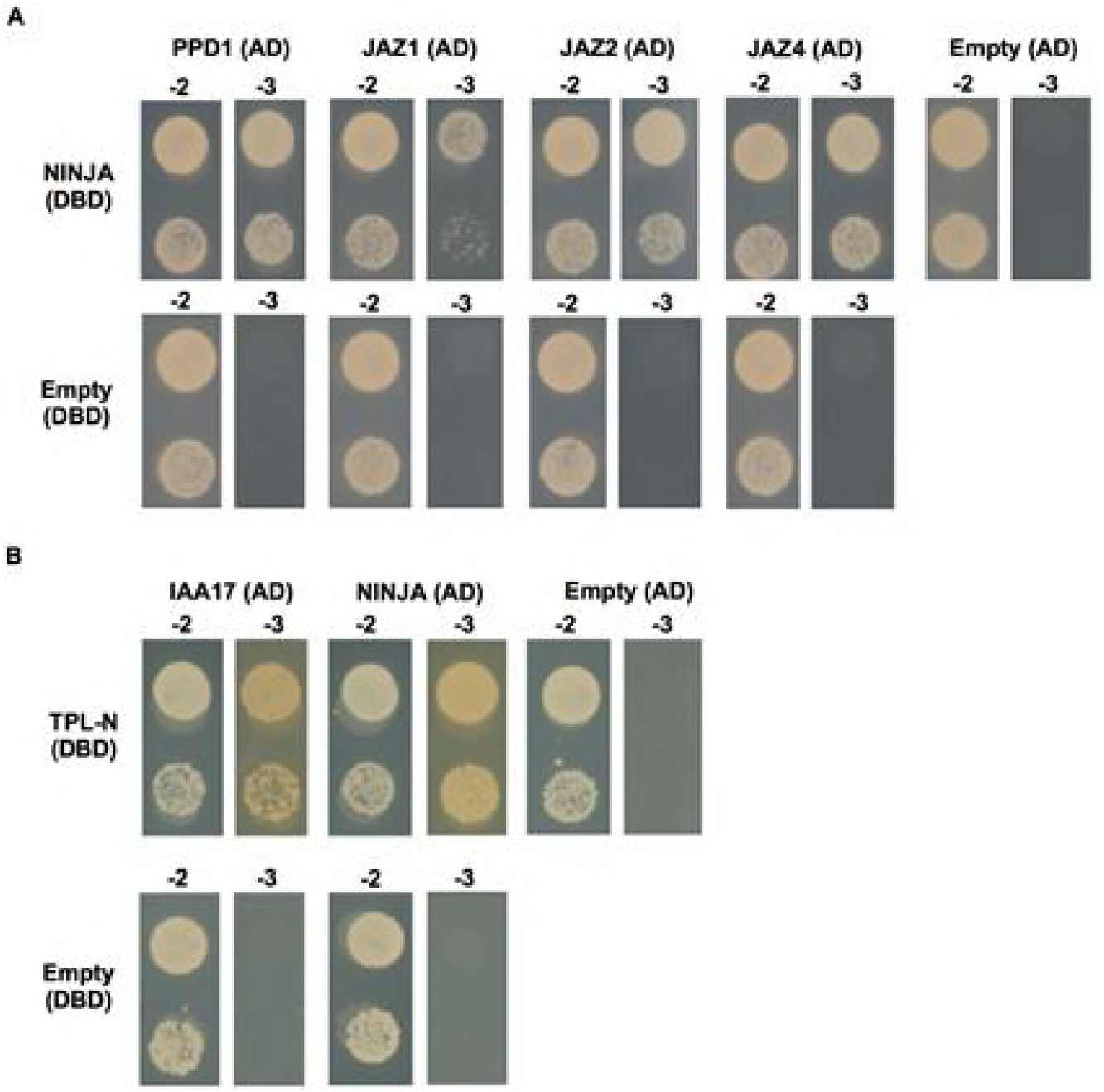
Y2H of the NINJA and TPL-N bait proteins with positive and negative control prey proteins. Y2H analysis of NINJA and TPL-N baits, fused to the DBD, and preys, fused to the AD, grown for 2 days on selective medium Synthetic Defined (SD)-Leu-Trp-His (−3). Transformed PJ69-4α yeast strains were also grown for 2 days on SD-Leu-Trp (−2) medium to confirm growth capacity. Direct interaction was confirmed between **(A)** NINJA and PPD1, JAZ1, JAZ2 and JAZ4, and **(B)** TPL-N and auxin/indole-3-acetic acid 17 (IAA17) and NINJA.

As expected, the binary interaction between the NINJA bait and the preys PPD1, JAZ1, JAZ2 and JAZ4 was confirmed (Fig 3A). Likewise, the TPL-N bait strain showed interaction with the preys auxin/indole-3-acetic acid 17 (IAA17) and NINJA (Fig 3B). Furthermore, neither of the bait strains exhibited auto-activation, which indicated that NINJA as well as TPL-N were functionally expressed in the bait strains.

#### Checkpoint 2: Evaluation of quality of Y2H-seq screening with bait strains by Sanger sequencing

For the actual Y2H screening, the bait strains were supertransformed with the *A. thaliana* Y2H cDNA prey library, followed by transformation efficiency assessment and five days of selective growth (Fig 4 and S1 Table). A minimum transformation efficiency of 1 x 10^6^ screened yeast colonies should be attained for a full Y2H cDNA library screening coverage. This benchmark was achieved for all Y2H screenings we performed (Table 1).

**Figure 4.**
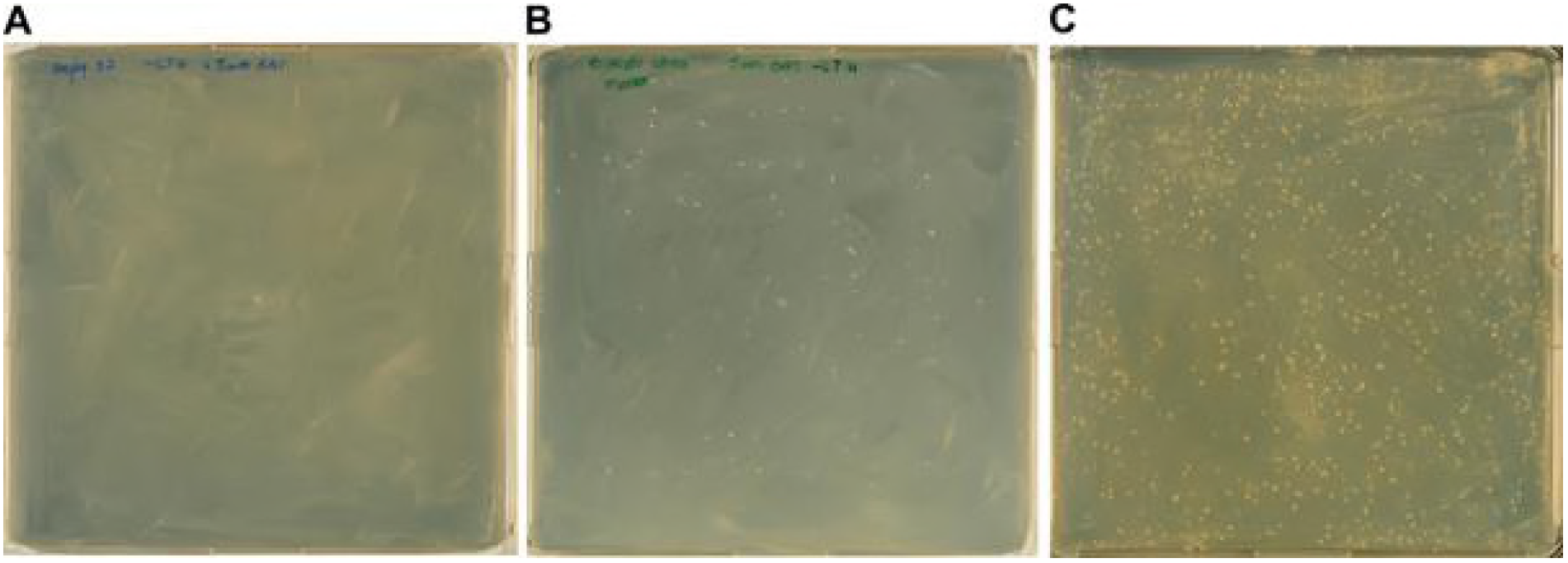
Y2H-seq selective growth. (A-C) The EMPTY (A), NINJA (B), and TPL-N (C) Y2H-seq screenings were performed on selective SD-Leu-Trp-His + 5mM 3-AT.

**Table 1.** Transformation efficiency of Y2H screenings using EMPTY, TPL-N and NINJA as baits. To ensure a full screening coverage of the *A. thaliana* Y2H cDNA library, screening of at least 1 x 10^6^ yeast colonies is advised [42].

| Bait | Transformation efficiency (# of colonies screened) |
| --- | --- |
| EMPTY | $1.23 \times 10^7$ |
| NINJA | $3.85 \times 10^6$ |
| TPL-N | $2.05 \times 10^7$ |

A minimum of ten individual colonies per screening were isolated, plasmids purified and the cDNA inserts of the prey plasmids Sanger-sequenced. In this second checkpoint, several known interactors could already be identified (Table 2). The ten sequences originating from the NINJA screening corresponded to two unique interaction partners that were previously described as NINJA interactors [17]. Likewise, the 12 prey sequences that corresponded to potential interactors of TPL-N were derived from six different, all known interactors [18].

**Table 2.** Sanger sequencing of isolated NINJA and TPL-N preys.

| # Colonies | Gene description | Gene ID |
| --- | --- | --- |
| **NINJA** |  |  |
| 8 | *A. thaliana* jasmonate-ZIM-domain protein 1 (JAZ1) | AT1G19180 |
| 1 | *A. thaliana* protein PEAPOD2 (PPD2) | ATt4G14720 |
| **TPL-N** |  |  |
| 4 | *A. thaliana* indole-3-acetic acid inducible 2 (IAA2) | AT3G23030 |
| 3 | *A. thaliana* indole-3-acetic acid inducible 28 (IAA28) | AT5G25890 |
| 2 | *A. thaliana* AGAMOUS-like 18 (AGL18) | AT3G57390 |
| 1 | *A. thaliana* indole-3-acetic acid inducible 4 (IAA4) | AT5G43700 |
| 1 | *A. thaliana* indole-3-acetic acid inducible 30 (IAA30) | AT3G62100 |
| 1 | *A. thaliana* indole-3-acetic acid inducible 9 (IAA9) | AT5G65670 |

#### Checkpoint 3: semi-quantitative qPCR, a complementary approach to evaluate the quality of a Y2H-seq screening

In a third checkpoint, the quality of the Y2H-seq screening was further assessed. All selectively grown yeast colonies were pooled per screening (Fig 2) and cDNA inserts of the prey plasmid pools were PCR-amplified with vector-specific primers (S2 Table). To examine whether potential interaction partners of the baits were overrepresented relative to the cDNA library control, a qPCR was performed using prey-specific qPCR primers (S2 Table). In the NINJA screen, compared to the control library, the genes encoding JAZ1, JAZ2, JAZ12, TIFY8 and PPD1 were overrepresented (Fig 5), in agreement with previous literature reports [17,34]. Hence, this shows the value of this qPCR assay set-up as a final checkpoint before the actual Y2H-seq analysis, at least for baits with a limited set of known interactors.

**Figure 5.**
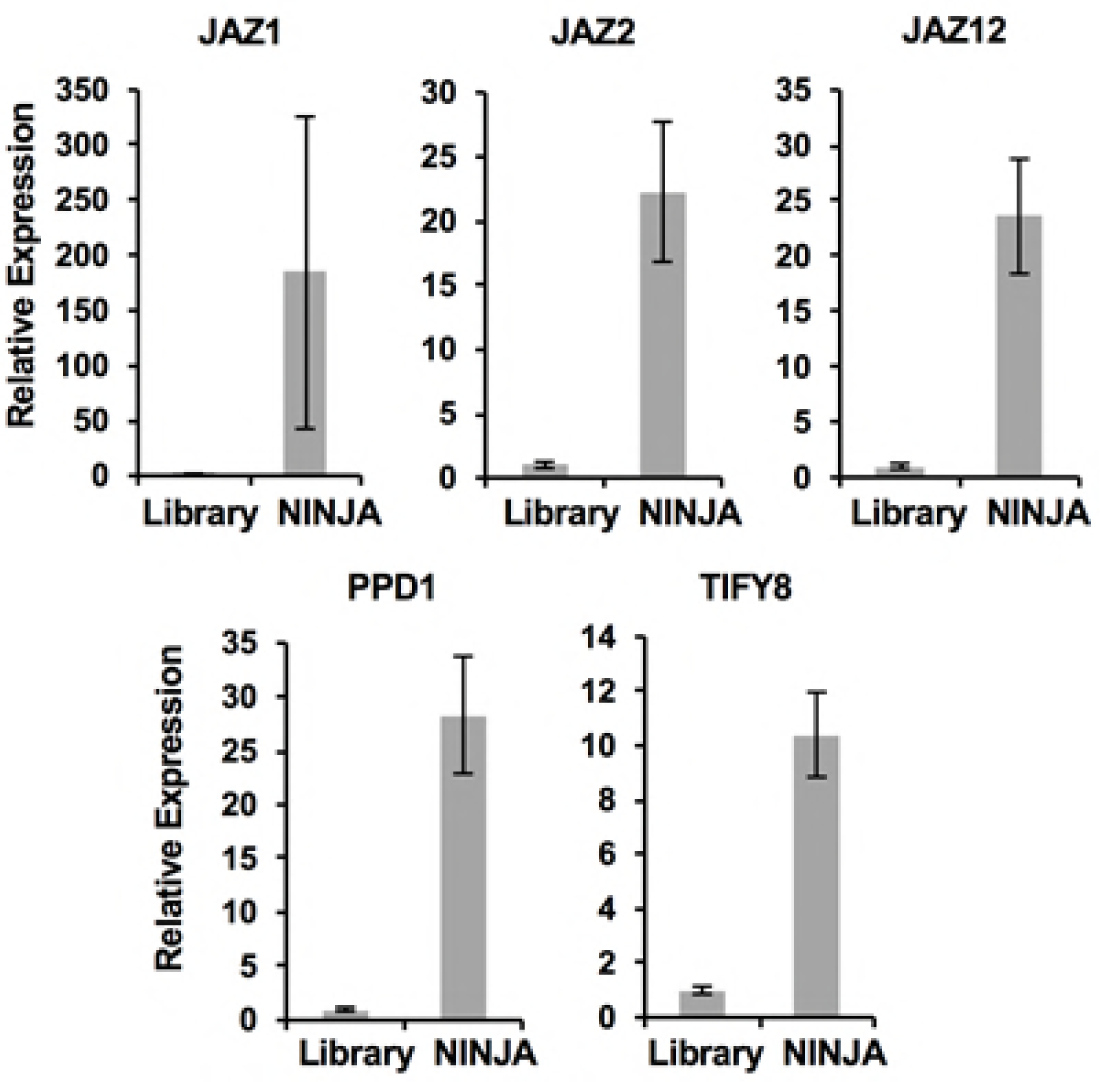
qPCR assessment of the NINJA Y2H-seq screen. *JAZ1*, *JAZ2*, *JAZ12*, *TIFY8* and *PPD1* were overrepresented in the PCR products of the NINJA screening compared to the PCR products of the *A. thaliana* cDNA library (Library).

In contrast to NINJA, TPL can interact with potentially hundreds of proteins [18]. Of the EAR-motif containing proteins known to interact with TPL and identified in the second checkpoint, only enrichment of IAA30 in the TPL-N pool could be observed (Fig 6, Table 2). Y2H cDNA library screenings are prone to false negatives, i.e. missing interactions, due among others to aberrant folding, clones with truncated genes or absence of the gene in the cDNA library. In the case of TPL-N, for example, the *NINJA* clone that is represented by the *A. thaliana* Y2H cDNA library was found to be truncated and missing the EAR domain necessary for binding with TPL-N. Therefore, critical analysis of theY2H cDNA library content prior and post Y2H screening remains crucial to critically interpret the outcome of a Y2H screen.

**Figure 6.**
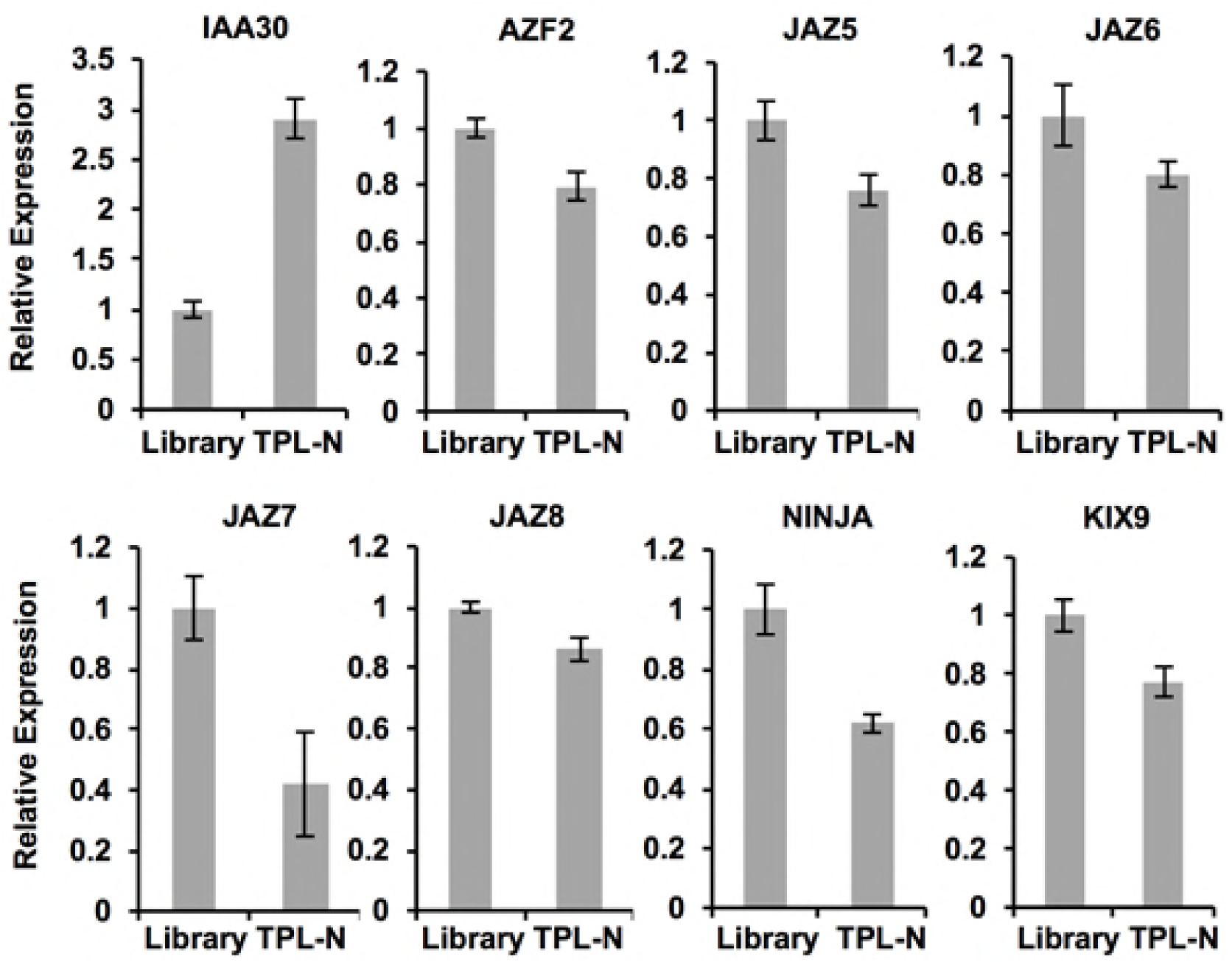
qPCR assessment of the TPL-N Y2H-seq screening. Only *IAA30* was overrepresented in the PCR products of the TPL-N screening compared to the *A. thaliana* cDNA library (Library) cDNA insert amplicons.

### Beyond the checkpoints: NGS of the amplified prey cDNA inserts

The prey pool amplicons of the EMPTY, NINJA, and TPL-N screenings were used as input for NGS by Illumina HiSeq 2000 (125-bp paired-end reads). Here, we used a pipeline relying on TopHat for read mapping and Cufflinks for gene expression quantification. The method presented here aims to compare the gene expression levels of the NINJA and TPL-N Y2H-seq screens with the EMPTY control screen to enrich for specific interaction partners while maximally avoiding the retrieval of false-positive interactions.

First, a quality check was performed on the raw reads. Thereby, adapters, low-quality sequences and partial vector sequences were trimmed. Concomitantly, paired-end and orphan single-end reads were split. The processed reads were then mapped to the reference genome (TAIR10) using TopHat. To avoid overestimation of short genes, only one mate-pair per read was used for mapping. The resulting alignments were used as input for Cufflinks, which generates the raw expression quantification data for each of the analyzed raw sequencing files. For the subsequent analysis of the raw expression data, a Y2H-seq pipeline was drafted in R-studio.

Mapped genes in the TPL-N and NINJA Y2H screenings with raw read counts less than six were eliminated. Genes in the EMPTY screening that had no raw read counts were given an arbitrary value of 1 and flagged as imputed. After calculating the Fragments Per Kilobase of Exon Per Million Fragments Mapped (FPKM) values, the signal to noise ratio (SNR) was defined for NINJA and TPL-N compared to EMPTY. Intuitively, one would expect little NGS data to be derived from the EMPTY screening, given that no yeast cells survived selective growth (Fig 4). However, this was not the case and can be explained by the pooling method employed here: ‘scraping’ all yeast cells from the selection plates includes also dead or nearly dead cells that may still contain intact prey plasmids. Hence, genes with a high representation in the cDNA library, and thus genes with a high expression level in *Arabidopsis* suspension cells, are identified in the EMPTY NGS data set.

Next, to allow setting relevant arbitrary thresholds, the 99.5^th^ percentiles of SNR_NINJA/EMPTY_ and SNR_TPL-N/EMPTY_ were calculated, leading to thresholds of 7.2 for NINJA and 6.0 for TPL-N screenings, respectively (Tables 3 and 4). With this first threshold, overall, from the 71 potential interactors of NINJA, seven were known to be interactors [17,34], whereas for TPL-N, 12 out of the 51 potential interactors had been previously reported [25].

**Table 3.**
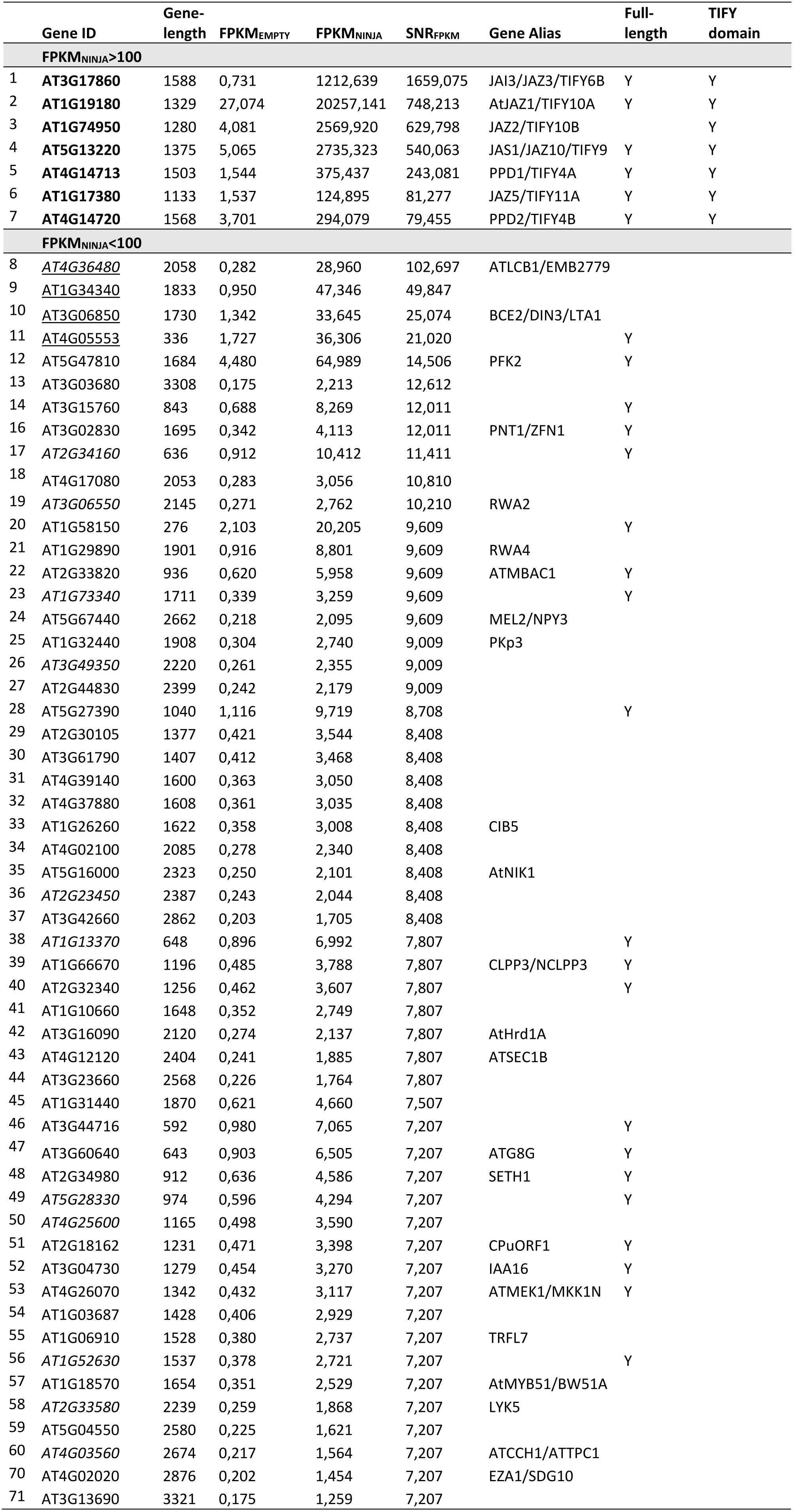
Signal-to-noise ratio of the FPKM values of NINJA and EMPTY Y2H-seq screenings. Genes with SNR_NINJA/EMPTY_>7.2 were retained, listed and ranked from high to low SNR. Flagged genes are italicized. Previously reported interactors of NINJA are indicated in bold. Potential interactors that were tested for binary interaction in further validation assays are underlined.

|  | Gene ID | Gene-length | FPKM<sub>EMPTY</sub> | FPKM<sub>NINJA</sub> | SNR<sub>FPKM</sub> | Gene Alias | Full-length | TIFY domain |
| --- | --- | --- | --- | --- | --- | --- | --- | --- |
|  | **FPKM<sub>NINJA</sub> > 100** |  |  |  |  |  |  |  |
| 1 | **AT3G17860** | 1588 | 0,731 | 1212,639 | 1659,075 | JAI3/JAZ3/TIFY6B | Y | Y |
| 2 | **AT1G19180** | 1329 | 27,074 | 20257,141 | 748,213 | AtJAZ1/TIFY10A | Y | Y |
| 3 | **AT1G74950** | 1280 | 4,081 | 2569,920 | 629,798 | JAZ2/TIFY10B |  | Y |
| 4 | **AT5G13220** | 1375 | 5,065 | 2735,323 | 540,063 | JAS1/JAZ10/TIFY9 | Y | Y |
| 5 | **AT4G14713** | 1503 | 1,544 | 375,437 | 243,081 | PPD1/TIFY4A | Y | Y |
| 6 | **AT1G17380** | 1133 | 1,537 | 124,895 | 81,277 | JAZ5/TIFY11A | Y | Y |
| 7 | **AT4G14720** | 1568 | 3,701 | 294,079 | 79,455 | PPD2/TIFY4B | Y | Y |
|  | **FPKM<sub>NINJA</sub> < 100** |  |  |  |  |  |  |  |
| 8 | <u>AT4G36480</u> | 2058 | 0,282 | 28,960 | 102,697 | ATLCB1/EMB2779 |  |  |
| 9 | <u>AT1G34340</u> | 1833 | 0,950 | 47,346 | 49,847 |  |  |  |
| 10 | <u>AT3G06850</u> | 1730 | 1,342 | 33,645 | 25,074 | BCE2/DIN3/LTA1 |  |  |
| 11 | <u>AT4G05553</u> | 336 | 1,727 | 36,306 | 21,020 |  | Y |  |
| 12 | AT5G47810 | 1684 | 4,480 | 64,989 | 14,506 | PFK2 | Y |  |
| 13 | AT3G03680 | 3308 | 0,175 | 2,213 | 12,612 |  |  |  |
| 14 | AT3G15760 | 843 | 0,688 | 8,269 | 12,011 |  | Y |  |
| 16 | AT3G02830 | 1695 | 0,342 | 4,113 | 12,011 | PNT1/ZFN1 | Y |  |
| 17 | <u>AT2G34160</u> | 636 | 0,912 | 10,412 | 11,411 |  | Y |  |
| 18 | AT4G17080 | 2053 | 0,283 | 3,056 | 10,810 |  |  |  |
| 19 | <u>AT3G06550</u> | 2145 | 0,271 | 2,762 | 10,210 | RWA2 |  |  |
| 20 | AT1G58150 | 276 | 2,103 | 20,205 | 9,609 |  | Y |  |
| 21 | AT1G29890 | 1901 | 0,916 | 8,801 | 9,609 | RWA4 |  |  |
| 22 | AT2G33820 | 936 | 0,620 | 5,958 | 9,609 | ATMBAC1 | Y |  |
| 23 | <u>AT1G73340</u> | 1711 | 0,339 | 3,259 | 9,609 |  | Y |  |
| 24 | AT5G67440 | 2662 | 0,218 | 2,095 | 9,609 | MEL2/NPY3 |  |  |
| 25 | AT1G32440 | 1908 | 0,304 | 2,740 | 9,009 | PKp3 |  |  |

|  |  |  |  |  |  |  |  |
| --- | --- | --- | --- | --- | --- | --- | --- |
| 26 | AT3G49350 | 2220 | 0,261 | 2,355 | 9,009 |  |  |
| 27 | AT2G44830 | 2399 | 0,242 | 2,179 | 9,009 |  |  |
| 28 | AT5G27390 | 1040 | 1,116 | 9,719 | 8,708 |  | Y |
| 29 | AT2G30105 | 1377 | 0,421 | 3,544 | 8,408 |  |  |
| 30 | AT3G61790 | 1407 | 0,412 | 3,468 | 8,408 |  |  |
| 31 | AT4G39140 | 1600 | 0,363 | 3,050 | 8,408 |  |  |
| 32 | AT4G37880 | 1608 | 0,361 | 3,035 | 8,408 |  |  |
| 33 | AT1G26260 | 1622 | 0,358 | 3,008 | 8,408 | CIB5 |  |
| 34 | AT4G02100 | 2085 | 0,278 | 2,340 | 8,408 |  |  |
| 35 | AT5G16000 | 2323 | 0,250 | 2,101 | 8,408 | AtNIK1 |  |
| 36 | AT2G23450 | 2387 | 0,243 | 2,044 | 8,408 |  |  |
| 37 | AT3G42660 | 2862 | 0,203 | 1,705 | 8,408 |  |  |
| 38 | AT1G13370 | 648 | 0,896 | 6,992 | 7,807 |  | Y |
| 39 | AT1G66670 | 1196 | 0,485 | 3,788 | 7,807 | CLPP3/NCLPP3 | Y |
| 40 | AT2G32340 | 1256 | 0,462 | 3,607 | 7,807 |  | Y |
| 41 | AT1G10660 | 1648 | 0,352 | 2,749 | 7,807 |  |  |
| 42 | AT3G16090 | 2120 | 0,274 | 2,137 | 7,807 | AtHrd1A |  |
| 43 | AT4G12120 | 2404 | 0,241 | 1,885 | 7,807 | ATSEC1B |  |
| 44 | AT3G23660 | 2568 | 0,226 | 1,764 | 7,807 |  |  |
| 45 | AT1G31440 | 1870 | 0,621 | 4,660 | 7,507 |  |  |
| 46 | AT3G44716 | 592 | 0,980 | 7,065 | 7,207 |  | Y |
| 47 | AT3G60640 | 643 | 0,903 | 6,505 | 7,207 | ATG8G | Y |
| 48 | AT2G34980 | 912 | 0,636 | 4,586 | 7,207 | SETH1 | Y |
| 49 | AT5G28330 | 974 | 0,596 | 4,294 | 7,207 |  | Y |
| 50 | AT4G25600 | 1165 | 0,498 | 3,590 | 7,207 |  |  |
| 51 | AT2G18162 | 1231 | 0,471 | 3,398 | 7,207 | CPuORF1 | Y |
| 52 | AT3G04730 | 1279 | 0,454 | 3,270 | 7,207 | IAA16 | Y |
| 53 | AT4G26070 | 1342 | 0,432 | 3,117 | 7,207 | ATMEK1/MKK1N | Y |
| 54 | AT1G03687 | 1428 | 0,406 | 2,929 | 7,207 |  |  |
| 55 | AT1G06910 | 1528 | 0,380 | 2,737 | 7,207 | TRFL7 |  |
| 56 | AT1G52630 | 1537 | 0,378 | 2,721 | 7,207 |  | Y |
| 57 | AT1G18570 | 1654 | 0,351 | 2,529 | 7,207 | AtMYB51/BW51A |  |
| 58 | AT2G33580 | 2239 | 0,259 | 1,868 | 7,207 | LYK5 |  |
| 59 | AT5G04550 | 2580 | 0,225 | 1,621 | 7,207 |  |  |
| 60 | AT4G03560 | 2674 | 0,217 | 1,564 | 7,207 | ATCCH1/ATTPC1 |  |
| 70 | AT4G02020 | 2876 | 0,202 | 1,454 | 7,207 | EZA1/SDG10 |  |
| 71 | AT3G13690 | 3321 | 0,175 | 1,259 | 7,207 |  |  |

**Table 4.** Signal–to-noise ratio of the FPKM values of TPL-N and EMPTY Y2H-seq screenings. Genes with SNR_TPL-N/EMPTY_>6 were retained, listed and ranked from high to low SNR. Flagged genes are italicized. Previously reported interactors of TPL are indicated in bold. Potential interactors that were tested for binary interaction in further validation assays are underlined. A ‘Y’ in bold font indicates the presence of an EAR domain in the wrong frame or in an untranslated region of the gene.

|  | Gene ID | Gene-length | FPKM<sub>EMPTY</sub> | FPKM<sub>N-TPL</sub> | SNR<sub>FPKM</sub> | Gene Alias | Full-length | EAR domain |
| --- | --- | --- | --- | --- | --- | --- | --- | --- |
|  | **FPKM<sub>N-TPL</sub> > 100** |  |  |  |  |  |  |  |
| 1 | **AT5G25890** | 873 | 11,301 | 3997,602 | 353,736 | IAA28/IAR2 | Y | Y |
| 2 | **AT3G23030** | 941 | 49,339 | 8581,581 | 173,933 | IAA2 | Y | Y |
| 3 | **AT5G43700** | 1168 | 0,994 | 111,604 | 112,307 | ATAUX2-11/IAA4 | Y | Y |
| 4 | **AT4G28640** | 1202 | 20,761 | 1121,873 | 54,037 | IAA11 | Y | Y |
| 5 | **AT4G29080** | 1337 | 4,341 | 219,498 | 50,568 | IAA27/PAP2 | Y | Y |
| 6 | **AT3G15540** | 970 | 7,180 | 266,613 | 37,135 | IAA19/MSG2 | Y | Y |
| 7 | **AT1G04250** | 1087 | 249,863 | 8692,277 | 34,788 | AXR3/IAA17 | Y | Y |
| 8 | <u>AT3G50000</u> | 1467 | 19,384 | 543,833 | 28,055 | ATCKA2 | Y |  |
| 9 | **AT2G33310** | 1820 | 64,093 | 1054,613 | 16,454 | IAA13 | Y | Y |
| 10 | AT2G46990 | 655 | 9,746 | 158,039 | 16,215 | IAA20 | Y | Y |
| 11 | **AT5G13790** | 962 | 15,685 | 198,181 | 12,635 | AGL15 | Y | Y |
| **FPKM<sub>N-TPL</sub> < 100** |  |  |  |  |  |  |  |  |
| 12 | <u>AT3G54390</u> | 1341 | 3,0294 | 69,6555 | 22,9933 |  | Y |  |
| 13 | **AT4G37940** | 715 | 0,8117 | 15,1114 | 18,6177 | AGL21 | Y |  |
| 14 | <u>AT2G40260</u> | 1233 | 0,4707 | 8,1976 | 17,4165 |  | Y |  |
| 16 | <u>AT3G58820</u> | 1391 | 0,4172 | 5,2619 | 12,6120 |  |  | **Y** |
| 17 | **AT1G51950** | 1539 | 4,9022 | 56,1646 | 11,4570 | IAA18 | Y | Y |
| 18 | **AT1G04100** | 1254 | 1,3884 | 15,8426 | 11,4108 | IAA10 | Y | Y |
| 19 | <u>AT3G05670</u> | 3090 | 0,9391 | 9,2492 | 9,8493 |  |  | Y |
| 20 | AT4G31620 | 1809 | 0,9624 | 9,2481 | 9,6091 |  |  | **Y** |
| 21 | <u>AT2G33550</u> | 1194 | 8,2629 | 77,0636 | 9,3265 |  |  | Y |
| 22 | AT1G08290 | 1760 | 0,3297 | 2,9705 | 9,0085 | WIP3 |  |  |
| 23 | <u>AT3G19860</u> | 1288 | 3,1540 | 25,7074 | 8,1506 | bHLH121 | Y |  |
| 24 | AT5G25160 | 959 | 1,2103 | 9,4494 | 7,8074 | ZFP3 | Y | Y |
| 25 | AT5G47110 | 1088 | 1,0668 | 8,3290 | 7,8074 | LIL3:2 |  |  |
| 26 | AT3G04730 | 1279 | 0,4537 | 3,5426 | 7,8074 | IAA16 |  | Y |
| 27 | AT5G04550 | 2580 | 0,2249 | 1,7562 | 7,8074 |  |  |  |
| 28 | AT3G15760 | 843 | 0,6884 | 4,9614 | 7,2068 |  | Y |  |
| 29 | AT1G12270 | 1949 | 0,5955 | 4,2919 | 7,2068 | Hop1 |  |  |
| 30 | AT3G47980 | 1097 | 0,5290 | 3,8126 | 7,2068 |  |  | **Y** |
| 31 | AT1G02650 | 1542 | 0,3764 | 2,7124 | 7,2068 |  |  |  |
| 32 | AT2G38950 | 2482 | 0,2338 | 1,6851 | 7,2068 |  |  |  |
| 33 | AT3G19070 | 1041 | 1,6725 | 11,7184 | 7,0066 |  |  | **Y** |
| 34 | AT1G28300 | 1317 | 0,8813 | 6,0868 | 6,9066 | AtLEC2 |  | **Y** |
| 35 | AT3G56250 | 669 | 0,8675 | 5,7308 | 6,6063 |  | Y |  |
| 36 | <u>AT1G01030</u> | 1905 | 0,3046 | 2,0126 | 6,6063 | NGA3 |  | **Y** |
| 37 | AT1G61900 | 1913 | 0,3034 | 2,0041 | 6,6063 |  |  | **Y** |
| 38 | AT1G79950 | 3123 | 0,1858 | 1,2276 | 6,6063 |  |  |  |
| 39 | AT3G43575 | 4332 | 0,1340 | 0,8850 | 6,6063 |  |  |  |
| 40 | AT5G36870 | 5616 | 0,2067 | 1,3033 | 6,3060 | ATGSL09/atgsl9 |  |  |
| 41 | AT2G30540 | 680 | 4,2672 | 26,6529 | 6,2459 |  |  |  |
| 42 | AT2G38110 | 1845 | 4,0891 | 25,5028 | 6,2367 | ATGPAT6 |  | **Y** |
| 43 | AT2G25180 | 1980 | 3,2241 | 19,8913 | 6,1695 | ARR12/AtARR12 |  | **Y** |
| 44 | <u>AT1G53030</u> | 530 | 1,0950 | 6,5762 | 6,0057 |  |  |  |
| 45 | <u>AT1G13680</u> | 1180 | 0,4918 | 2,9537 | 6,0057 |  |  |  |
| 46 | <u>AT4G19540</u> | 1215 | 0,4776 | 2,8686 | 6,0057 | INDH/INDL | Y |  |
| 47 | AT3G56160 | 1600 | 0,3627 | 2,1784 | 6,0057 |  |  |  |
| 48 | <u>AT5G03570</u> | 1673 | 0,3469 | 2,0833 | 6,0057 | ATIREG2/FPN2 |  |  |
| 49 | AT3G59150 | 1866 | 0,3110 | 1,8678 | 6,0057 |  | Y |  |
| 50 | <u>AT4G03560</u> | 2674 | 0,2170 | 1,3034 | 6,0057 | ATCCH1/ATTPC1/FOU2 |  |  |
| 51 | AT3G03680 | 3308 | 0,1754 | 1,0536 | 6,0057 |  |  |  |

When super-implying a second threshold, in this case of >100 on the FPKM_NINJA_ and FPKM_TPL-N_ values, nearly all retained interactors were either reported already or very plausible. Indeed, in the case of NINJA, only TIFY-domain containing proteins were retained (Fig 6, Table 3). In the case of TPL-N, all but one of the retained proteins using this second threshold contained an EAR-motif [43], the conventional TPL recruitment domain (Fig 7, Table 4), and also includes proteins not yet individually reported as TPL-interactors, but belonging to multigene families such as the AGAMOUS-LIKE (AGL) and INDOLE-3-ACETIC ACID INDUCIBLE (IAA) proteins, many members of which have already been reported as TPL interactors [18,25]. Together, this demonstrates the robustness and potential of the designed Y2H-seq platform.

**Figure 6.**
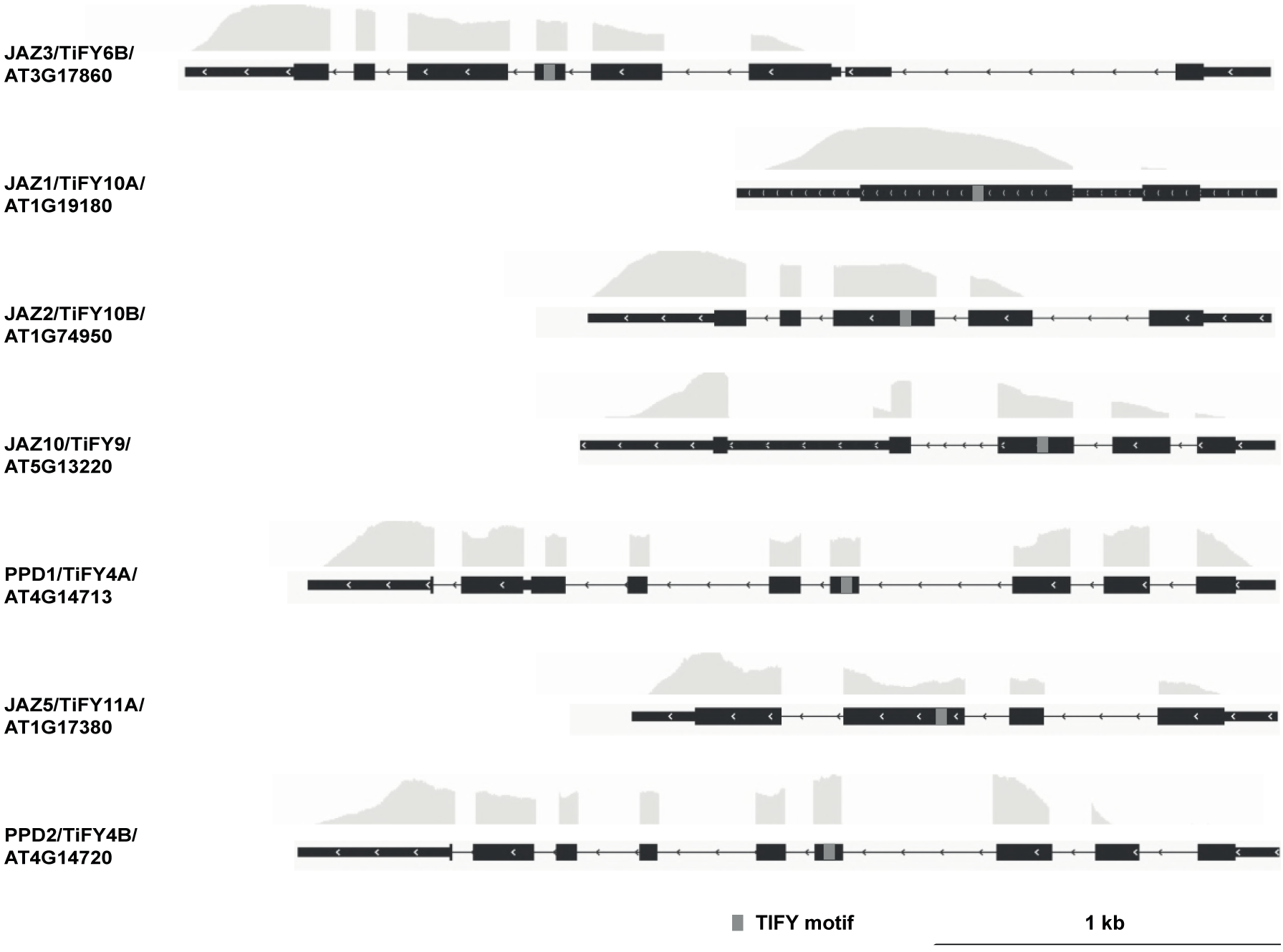
Deepseq coverage of the NINJA interactors using cutoff of SNR_NINJA/EMPTY_>7.2 and FPKM_NINJA_>100. The depth of the deepseq coverage for each gene, visualized by the coverage track, is aligned to the gene model. Coding sequences are represented by thick black boxes, 5’ and 3’ untranslated regions by thin black boxes and introns by thin black lines, respectively. The light grey boxes in the gene model correspond to the TIFY motif.

**Figure 7.**
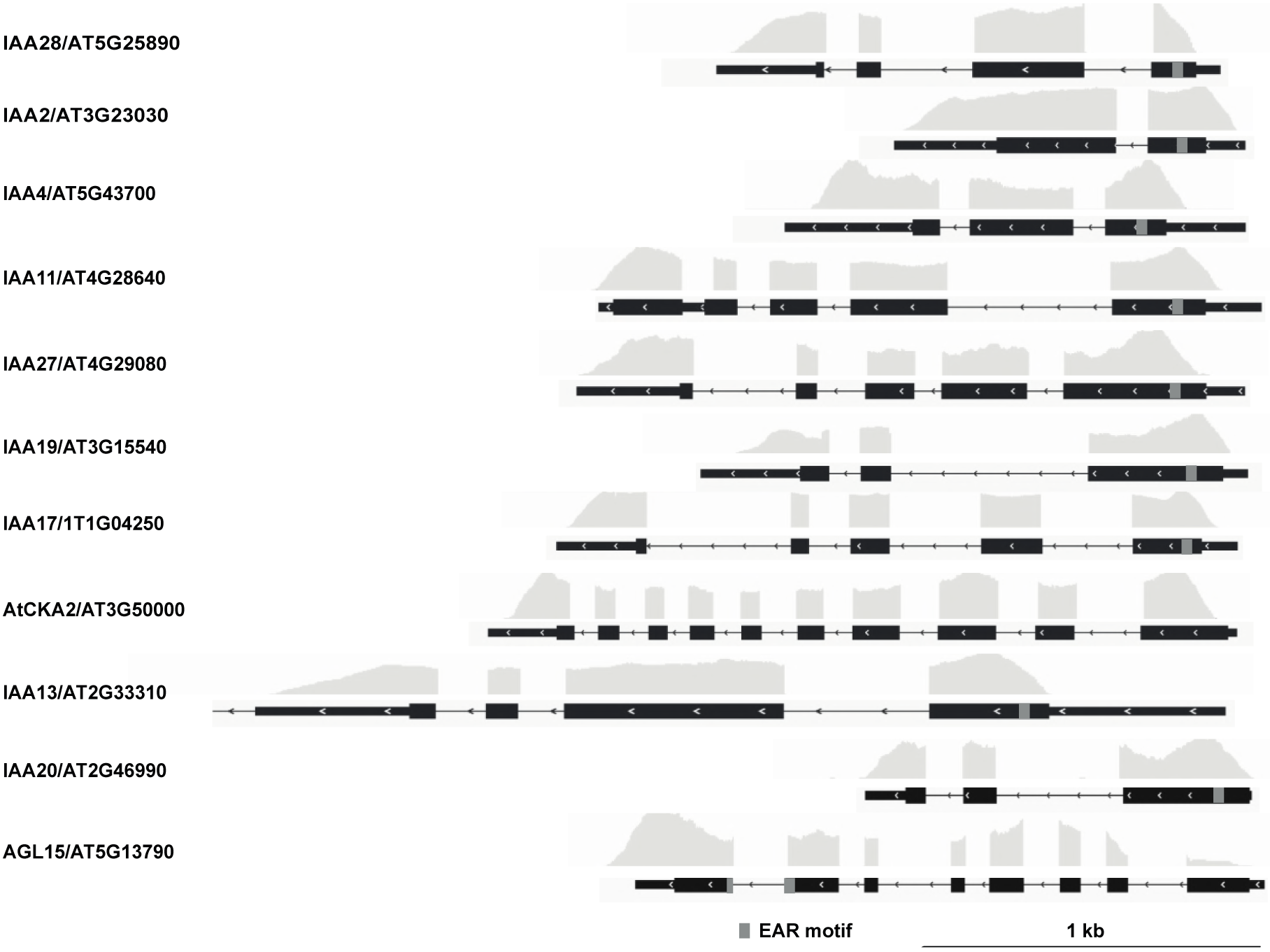
Deepseq coverage of the N-TPL interactors using cutoff of SNR_N-TPL/EMPTY_>6 and FPKM_N-TPL>_100. The depth of the deepseq coverage for each gene, visualized by the coverage track, is aligned to the gene model. Coding sequences are represented by thick black boxes, 5’ and 3’ untranslated regions by thin black boxes and introns by thin black lines, respectively. The grey boxes in the gene model correspond to the EAR motif.

To assess whether the retrieved preys that did not pass our stringent cut-offs, nonetheless represent true potential interactors of NINJA and N-TPL, additional Y2H experiments were carried out. For NINJA, the first four potential interaction partners with SNR_NINJA/EMPTY_>7.2 and FPKM_NINJA_<100 were tested in a binary Y2H assay (Table 3 and Figure 8). However, none of them showed interaction with NINJA, indicating that the installed threshold of SNR_NINJA/EMPTY_>7.2 and FPKM_NINJA_ >100 served as a good selection criterion, at least for NINJA.

**Figure 8.**
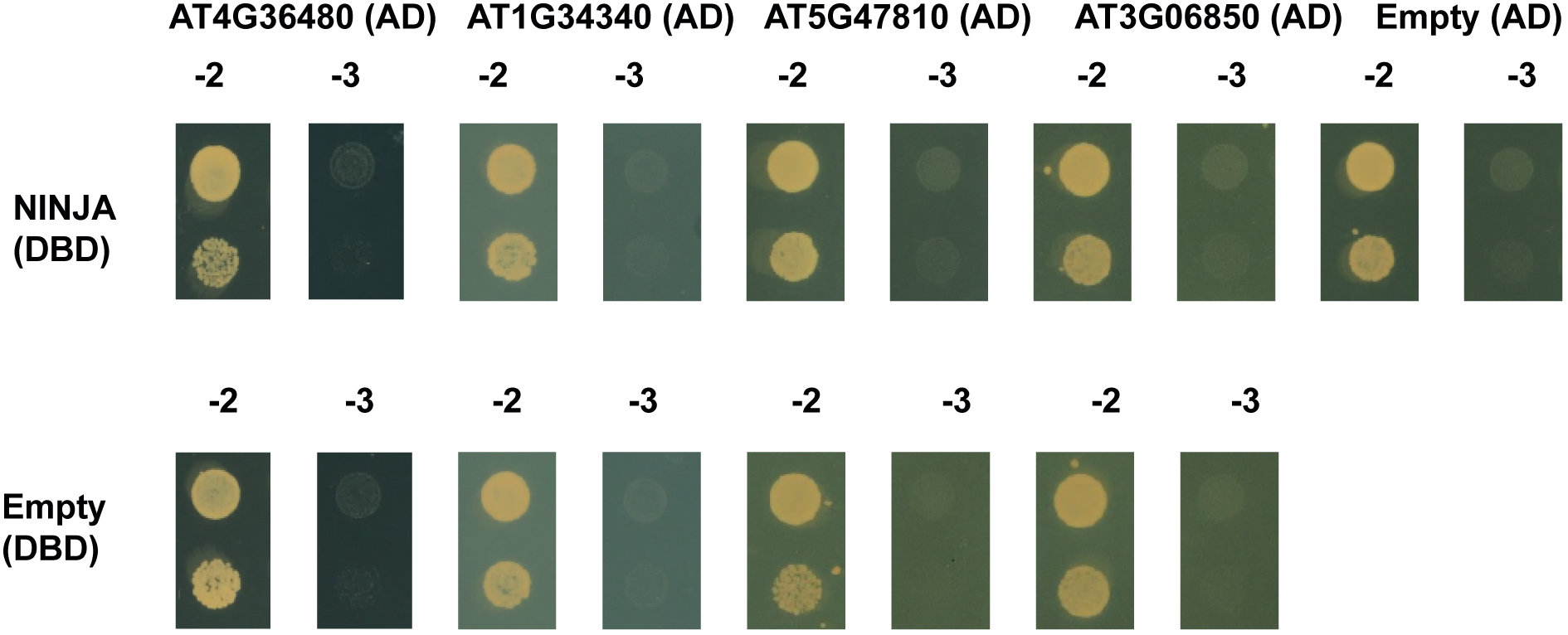
Y2H analysis of potential interaction partners of NINJA. Y2H analysis of NINJA, fused to the DBD, and potential interaction partners, fused to the AD of the GAL4 TF, grown on selective medium SD-Leu-Trp-His (−3). Transformed PJ69-4α yeast strains were also grown on SD-Leu-Trp (−2) medium confirm growth capacity. No direct interactions could be observed for retrieved preys below the threshold of SNR_NINJA/EMPTY_>7.2 and FPKM_NINJA_ values>100. Interaction between TPL and AT2G33550 had previously also been detected in the TOPLESS Interactome Y2H screen [18].

In the retained list of potential interactors using threshold SNR_TPL-N/EMPTY_>6 with FPKM_N-TPL_>100 values, the one candidate ATCKA2 (AT3G50000) that did not contain an EAR-domain was tested for direct interaction with N-TPL in a Y2H assay, besides five candidates with FPKM_N-TPL_<100 (Table 4 and Figure 9). For the latter set, we specifically avoided to pick candidates from the AGL and IAA families, which are most likely true, but less abundant interactors, and chose both candidates with and without an EAR domain. ATCKA2 interaction with N-TPL could not be confirmed with binary Y2H, suggesting it was a false positive caused by the Y2H-seq pipeline. In contrast however, interaction between TPL-N and the five other candidates were all confirmed, demonstrating that they do not represent artefacts of the Y2H-seq methodology and may be true interactors. Hence, in contrast to NINJA, this implicates that the arbitrary threshold of SNR_TPL-N/EMPTY_>6 with FPKM_N-TPL_>100 was too stringent for N-TPL. Perhaps this may be due to the pleiotropic function of TPL, which has an exceptionally high number of protein interactors, often from multigene families. For proteins such as NINJA, with a more defined role and a well-defined set of interactors, a stricter threshold may be justified. For proteins such as TPL, one may need to be more relaxed in determining candidate interactors. As exemplified here, this leads to the identification of potential novel interactors from gene families previously unreported to be capable of interacting with TPL, including EAR-domain containing proteins such as the RING/U-box protein AT3G05670, or proteins that do not contain an EAR domain such as the putative TF AT3G54390, the homeodomain TF AT2G40260 and the bHLH TF AT3G19860 (Table 4 and Figure 9).

**Figure 9.**
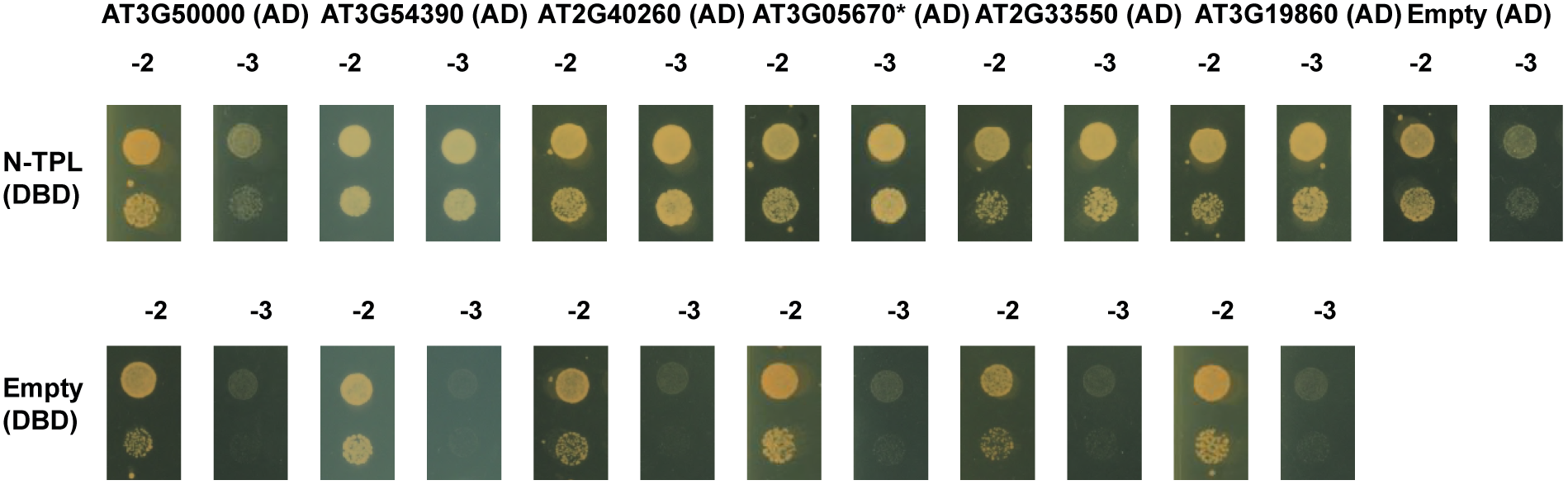
Binary Y2H validation of potential interaction partners of N-TPL. Y2H analysis of N-TPL, fused to the DBD, and potential interaction partners, fused to the AD of the GAL4 TF, grown on selective medium SD-Leu-Trp-His (−3). Co-transformed PJ69-4α yeast strains were also grown on SD-Leu-Trp (−2) medium to confirm growth capacity. No direct interaction was confirmed between ATCKA2 encoded by AT3G5000 and N-TPL, in contrast to the interactions with all other potential interactors selected from the list with a threshold of SNR_NINJA/EMPTY_>7.2 and FPKM_NINJA_<100 values. * indicates a truncated version of the protein, as it was present in the Y2H cDNA library.

## Discussion

Here, we present a newly designed high-throughput Y2H-seq strategy to identify PPIs, which enables exploiting the full qualitative and quantitative potential of Y2H library screenings in an unprecedented way. Our method circumvents multiple shortcomings of a conventional Y2H library screening. As such, for instance consumable and DNA sequencing costs are significantly cut by using a pool-based NGS-strategy instead of the conventional isolation, manipulation and sequencing of individual yeast clones that survive the screening selection. Moreover, a higher sensitivity can be achieved in our Y2H-seq strategy through maximal coverage of PPIs by increasing library titers. Consequently, interactions with less abundant proteins that would be masked or lost in conventional Y2H screenings can now be detected. In this regard, a factor that will determine the impact of future Y2H-seq screenings more than ever, will be the choice and the quality of the Y2H cDNA library. For instance, full-length protein libraries may mask PPIs by steric hindrance, hence the use of more complex Y2H cDNA libraries encoding protein fragments as well as full-length proteins may now be considered, and screened in one effort, which could lead to a comprehensive coverage of the PPI space. The utility of fragment-based Y2H approaches has previously been demonstrated [44,45]. By playing with sample preparations to generate cDNA libraries, one could increase the genome coverage with no extra effort in the Y2H screening. For instance, different organs from a single plant, different developmental stages of a single organ, or explants subjected to different environmental cues or chemicals can now be pooled in a single cDNA library. This will allow expanding the number of genes screened in a single event, as well as different versions of the same gene, e.g. following expression after alternative splicing or translation start events. As such, the Y2H-seq strategy will provide an effective way to discover differentially regulated PPIs, allowing further exploration of biological pathways and their regulation. Furthermore, the use of cDNA libraries makes it possible to identify novel interaction partners of organisms of which the genome has not been fully annotated yet, unlike the use of ORF libraries based on known and completely fixed gene models.

The Y2H-seq strategy implements a quantitative readout system, with a straightforward and adaptable scoring procedure. The use of background controls reliably allows eliminating false positives in early stage. This does not only involve comparing quantitative NGS readouts from Y2H-seq screenings with bait proteins to those of control screenings with ‘empty’ control vectors, but also comparing the readouts of the screenings with bait proteins among each other. Indeed, as is also the case with other PPI discovery methods, such as tandem affinity purification [46,47], a specific ‘blacklist’ of returning Y2H-seq interactors for each cDNA library can be composed by marking common interactors of seemingly unrelated bait proteins. This may allow fine-tuning the thresholds to be set up in the filtering of the Y2H-seq NGS data, and thereby enable determining robust priority lists and reducing laborious and needless downstream validation assays to a minimum.

Finally, this strategy can also easily be extended to Y1H screenings, for which the same cDNA library could be screened, but for which considerably higher false-positive rates are typically obtained as compared to Y2H screenings [48,49]. As such, we anticipate that the cost and labor reduction along with the increased detection and quantification potential of our Y2H-seq strategy can give an important upgrade to this long-existing, but far from fully exploited screening tool.

## Acknowledgements

We thank Annick Bleys for helping to prepare the manuscript and Frederik Coppens for helpful advice on NGS. This research was supported by the Research Foundation Flanders for postdoctoral fellowships to J.P. and L.P., the Program Ciências Sem Fronteiras for a predoctoral fellowship to B.R. (Grant 201135/2014-0), the BEC.AR program for overseas training of Argentine professionals in the fields of science, technology and productive innovation for a scholarship to M.P. and the Special Research Fund from Ghent University (project O1J14813).

## Supporting information

**S1 Table.**
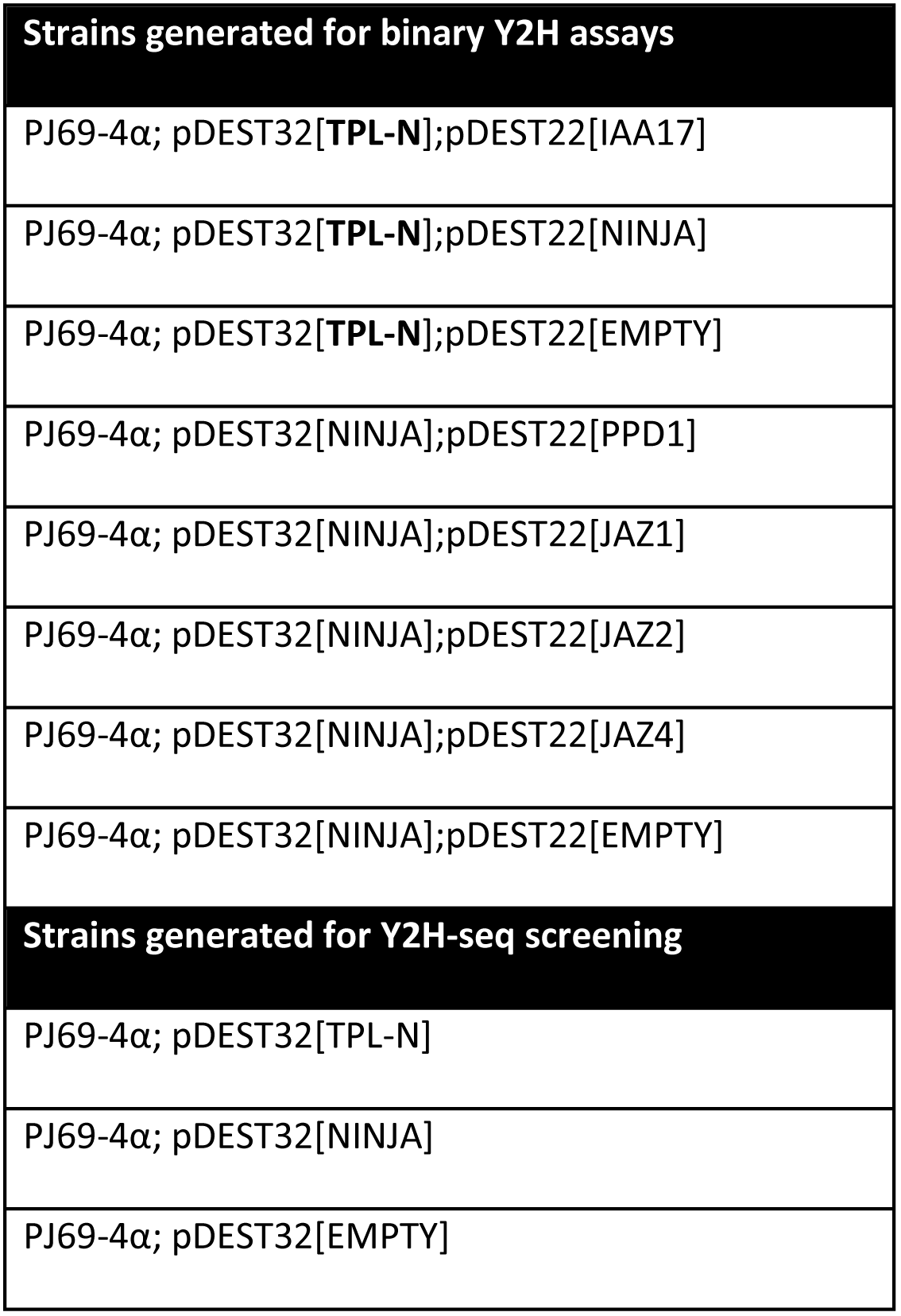
Yeast strains generated in this study.

**S2 Table.**
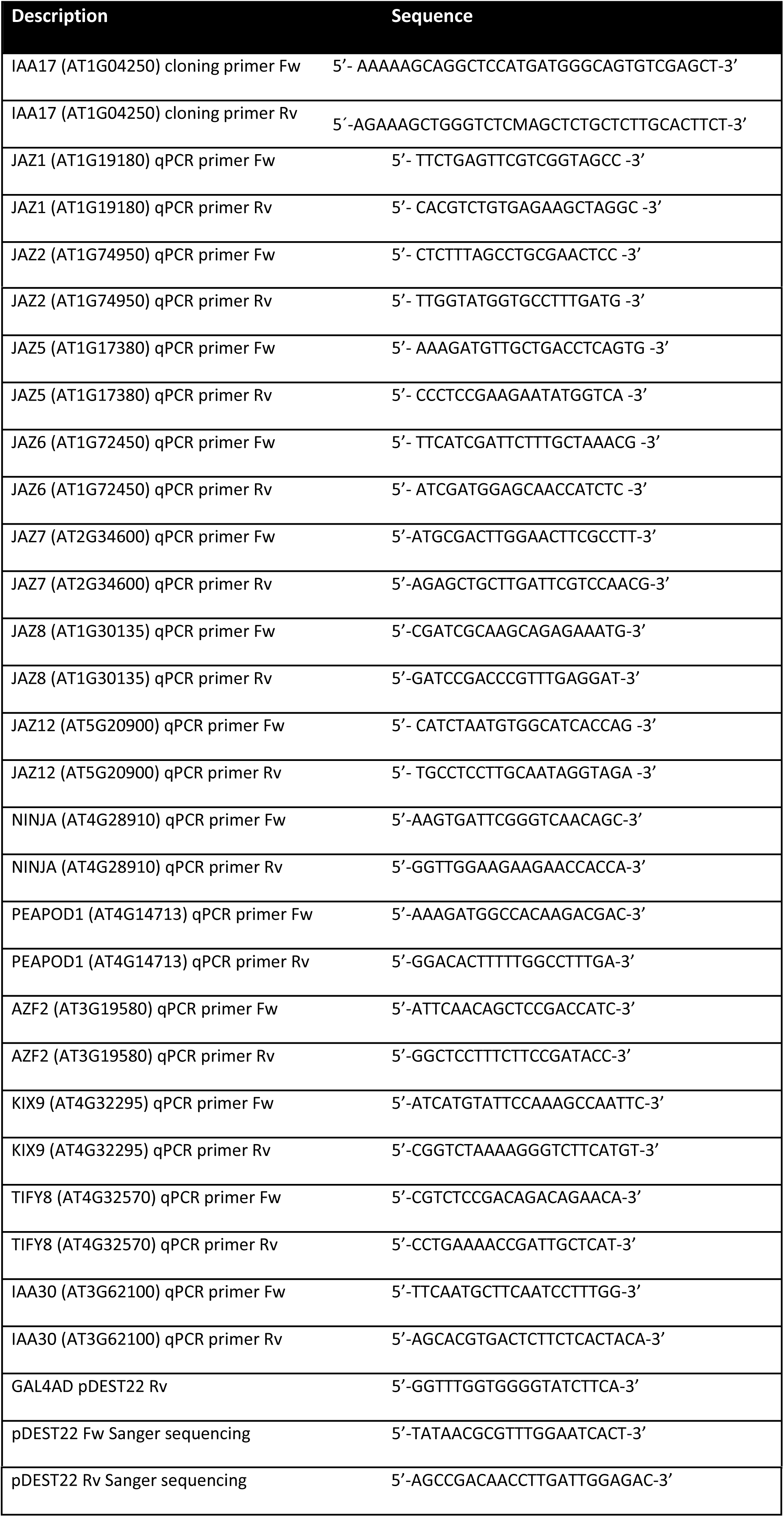
Primers used in this study.

